# Migration and the excess exposure of birds to human density in North America

**DOI:** 10.1101/2022.11.28.518244

**Authors:** Erin K. Jackson, Roslyn Dakin

## Abstract

Migratory species must cross a range of landscapes that are increasingly modified by humans. A key question is how migrating populations are responding to human-induced environmental change. Here, we model the spring migration dynamics of 63 bird species in North America to quantify their exposure to human population density. We find that most bird species have a negative navigational bias, suggesting that they attempt to avoid human-dense areas during migration, and yet they experience far greater human density during migration as compared to breeding. Species that experience excess human density during migration share several key traits: they tend to be nocturnal migrants, they start migrating through North America earlier in the year, and they tend to migrate longer distances. These findings underscore that birds are especially vulnerable to threats associated with human disturbance during migration, with predictable exposures that are often elevated by 2- to 3-fold during migration.

## 1. INTRODUCTION

During spring migration, hundreds of bird species travel across a broad range of landscapes to their breeding grounds. Migrants are faced with key physiological and behavioural challenges, including human-imposed obstacles such as artificial light, window collisions, habitat loss and fragmentation, as well as altered food availability and foraging periods (Isaksson 2018). These challenges may have important impacts on the ability of migrants to successfully reach their breeding grounds. For example, window collisions and domestic cats are two major sources of mortality for migrating birds (Loss *et al*. 2013, 2014; Machtans *et al*. 2013), artificial light fragments the aerial landscape and impairs navigation (Van Doren *et al*. 2017; Korpach *et al*. 2022), and loss of stopover areas can limit access to resources necessary to fuel migration flight. Although most studies of the impact of human activity on birds have been conducted during the breeding phase (Faaborg *et al*. 2010), recent work reveals that some bird species occur in more human-modified sites during migration as compared to other phases of the annual cycle (Zuckerberg *et al*. 2016; Cabrera-Cruz *et al*. 2018). This raises key questions as to whether and how migrating species differ in their response to human activity (Zuckerberg *et al*. 2016; Cabrera-Cruz *et al*. 2018).

Here, we study the spring migration dynamics of 63 North American landbird species to investigate how migration is influenced by human population density (HPOP). We focus on HPOP because it captures a suite of human footprint effects on the landscape, including building density, the density of roads and impervious surfaces, changes in resource availability, light and noise pollution, and chemical contaminants, among other factors. We expect bird species to differ in their responses to HPOP through multiple causal pathways. Importantly, our aim in this study is not to disentangle how different mechanisms occur across species, but rather to quantify how migrating bird species are responding to human density in general, and to investigate sources of variation in their exposure. To model migration dynamics, we use data from eBird, a community science database based on checklists submitted by hundreds of thousands of regular users (Sullivan *et al*. 2009; Strimas-Mackey *et al*. 2020). These widespread observations make eBird particularly useful for modelling the timing and trajectory of migratory populations (Fink *et al*. 2011; La Sorte *et al*. 2013, 2016; Feng *et al*. 2021; La Sorte & Horton 2021). eBird checklists also cover a wide range of HPOP in North America, with the greatest representation covering sites that range between 10-10,000 persons/km^2^ (e.g., Figure 1A inset; see also supplement Figure S1).

**Figure 1.**
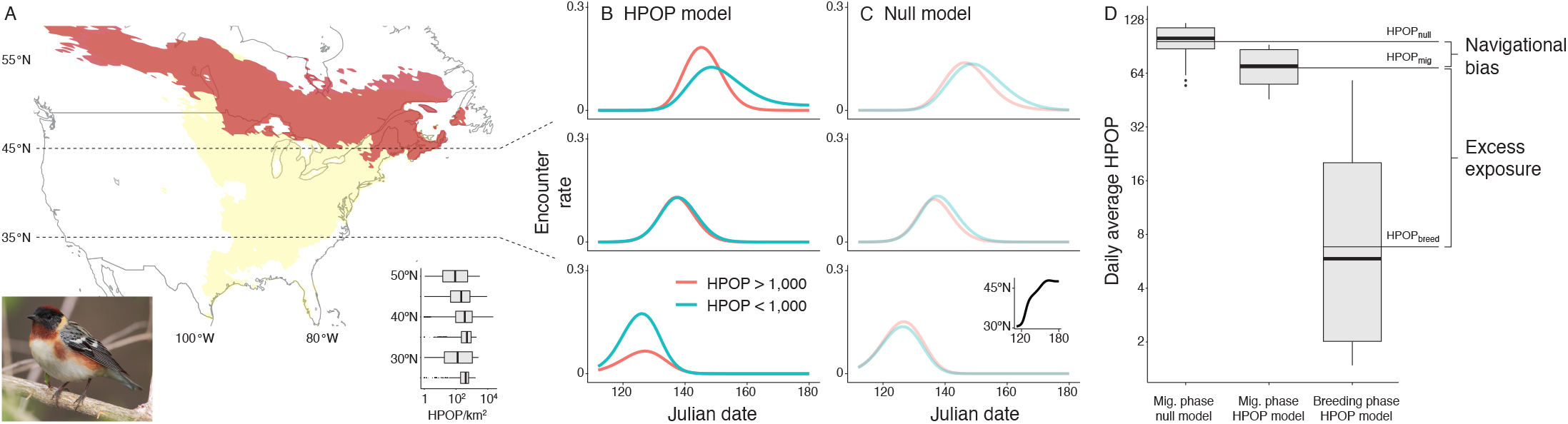
Modeling spring migration dynamics. An example of the modelling procedure is shown here for the bay-breasted warbler (*Setophaga fusca*). A) North American range map for *S. fusca*. The darker shaded regions represent the species’ breeding range, and the lighter shaded regions represent its migration range, as determined by the eBird Status and Trends Project. The inset shows the distribution of HPOP for eBird checklists in the species’ 2019 analysis. Notably, the range of HPOP characteristics is broadly similar across latitudes. B) Fitted GAM of the bay-breasted warbler’s spring migration in 2019. Each row shows encounter rates averaged over three latitude bands (below 35°N; 35-45°N; and above 45°N, as shown via the horizontal dotted lines), and two HPOP categories (below vs. above 1,000 persons/km^2^). Note that latitude and HPOP are categorized here to allow visualization of the complex model fit, but the analysis treats these variables as continuous. C) Null model of bay-breasted warbler spring migration. The inset shows the species’ estimated latitudinal progress. D) Daily HPOP estimates for the bay-breasted warbler in 2019. Mean values were used to calculate metrics of navigational bias and excess exposure. Photo by Wikimedia Commons user Mdf.

Our first aim in this study is to estimate average levels of HPOP experienced during the spring migration phase and breeding phase, respectively (hereafter, we refer to these estimates as HPOP_mig_ and HPOP_breed_). We also examine how these estimates relate to species differences in breeding SSI (Julliard *et al*. 2006), an index of how specialized a species is in terms of its breeding habitat. We predicted that more specialized species with a higher breeding SSI would have lower HPOP_mig_ and HPOP_breed_, because narrow resource requirements are less likely to be met in urbanized areas (Bonier *et al*. 2007; Ducatez *et al*. 2018; Callaghan *et al*. 2019). Second, following recent work establishing that migrating populations often use regions that differ in landcover and light pollution (Zuckerberg *et al*. 2016; Cabrera-Cruz *et al*. 2018), we calculate the average excess HPOP experienced by each species during migration as compared to breeding. We also estimate each species’ navigational bias with respect to HPOP, which we define as the difference between HPOP_mig_ and a null expectation. Finally, we use a comparative analysis to examine species differences in exposure to excess HPOP during migration, which we define as the difference between HPOP_mig_ and HPOP_breed_. We consider as predictors several traits related to migration biology, including the annual timing of a species’ migration through North America, the distance the species migrates, and the time of day (nocturnal or diurnal) when migration flight primarily occurs (Gauthreaux & Belser 2006; Van Doren *et al*. 2017; McLaren *et al*. 2018; Adams *et al*. 2021). Overall, this is the first work to characterize how a broad range of migratory bird species respond to HPOP during migration.

## 2. MATERIALS AND METHODS

### 2.1 General approach

We selected 63 migratory terrestrial bird species that are common in North America, that overwinter primarily at latitudes below 40°N, and that breed primarily at latitudes above 45°N. Given that migration behaviour can vary from year-to-year, we chose to study spring migration in three successive years, 2017-19. To model migration dynamics for each species-year, we used observations from eBird, fitting a generalized additive model (GAM) for each species-year. Then, we extracted downstream estimates from the fitted species-year models to characterize a set of responses to HPOP. All data preparation and analyses were performed in R 3.6.3 (R Core Team 2020). Further details of the study species are provided in the supplement and Table S1.

### 2.2 eBird and HPOP data

eBird data are provided in birdwatching checklists, which contain a record of the types and counts of species observed by an eBird participant. We downloaded the eBird Sampling Event Dataset for the years 2017, 2018 and 2019, which includes meta-data for all checklists in those years. As per the guidelines outlined by eBird (Johnston *et al*. 2021), we included only ‘complete’ checklists that were classified as either ‘Stationary’ or ‘Traveling’, with a duration less than 300 minutes and a distance less than 5 km. ‘Complete’ checklists are those where all species observed were reported, which is important for inferring species absences (Strimas-Mackey *et al*. 2020). We limited our analysis to checklists that started between 04:00 and 20:00 local time with ten or fewer observers. If multiple observers submitted checklists for the same event, we retained only one checklist. For the years we examined, 80% of checklists were classified as complete, and of those, 98% were stationary or traveling (average duration of 65.6 minutes, average distance of 3.35 km).

We downloaded occurrence data for each species-year from its eBird Basic Dataset in the USA and Canada for the period from January 1 to June 30. This period encompasses spring migration as well as the pre- and post-migration periods, to ensure that we could characterize the full spring migration phase for all study species. We designated sites as latitude and longitude coordinates gridded to 0.1° and defined each species-year ‘range’ as sites with at least one observation of that species. We then compiled a species-year dataset based on checklists that occurred at those sites from January 1 to June 30, and removed sites that had fewer than 50 checklists from further analysis for a given species-year. The species-year datasets had an average of 0.84 million checklists (+/- SD 0.43 million), with the smallest sample size > 96,000 checklists.

To determine HPOP values, we used data from the Gridded Population of the World (GPW), Version 4: Population Density, Revision 11 for the year 2015 (CIESIN 2016). GPW uses census data to model global human population densities (persons/km^2^) at high resolution (30 arcseconds, ∼1 km at the equator). Using the raster package v. 3.4-10 (Hijmans 2021), we extracted HPOP for each unique coordinate pair in the eBird Sampling Event Dataset, gridded to the nearest 0.01°, as the mean natural log-transformed human population density for a 5 km radius around those coordinates. The 5 km radius accounts for the relative mobility of birds, as well as previous work showing that artificial light affects birds primarily within this spatial scale (Van Doren *et al*. 2017; McLaren *et al*. 2018).

### 2.3 Modelling spring migration dynamics

Spring migration generates a pulse in occurrence at a given location as birds arrive, and the timing of this pulse depends on latitude as well as other site characteristics (Figure 1A-B). We used GAMs to model these nonlinear dynamics. GAMs are useful for this purpose because they sum together a series of smooth functions that allow for changes in the nonlinear landscape for each cross-section of the data. We fit a separate binomial GAM for each species-year dataset using the mgcv package v. 1.8-31 (Wood 2011), with the response variable as the binary detection of a species as a function of the following predictors:

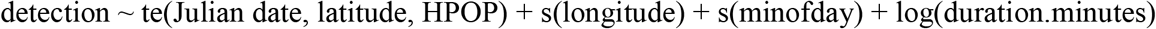

The te function (tensor product smooth) models nonlinear interactions among terms with different units of measurement (Wood *et al*. 2013), which is important because we expect the effect of HPOP on occurrence to depend on latitude and date. This is illustrated in Figure 1A-B showing an example for the bay-breasted warbler. The fitted model describes how the pulse of warblers occurs first at lower latitudes, and later at higher latitudes. In addition, the fitted model predicts that at lower latitudes in the month of April, this particular warbler species is detected much more frequently in low HPOP sites as compared to high HPOP sites (Figure 1B). By late

May, the model predicts that this species is detected at high-latitude, high-HPOP sites prior to lower HPOP areas at those latitudes (Figure 1B). These patterns differ across species; as an additional example, we show the same plots for the cedar waxwing in the supplement (Figure S2).

The GAMs also included the time of day when the checklist began (in minutes) and the checklist duration (in minutes, log-transformed) because these two variables are known to influence detection probability for eBird observers (Strimas-Mackey *et al*. 2020; Johnston *et al*. 2021). We describe the predictions of the fitted GAMs as ‘encounter rates’, or the predicted probability that an observer will report a given species for a standardized set of observation or effort parameters (Strimas-Mackey *et al*. 2020). These encounter rate values are used for subsequent estimates of HPOP_mig_ and HPOP_breed_, which are described in the sections below.

To check the predictive accuracy of the fitted species-year GAMs, we used cross-validation by first fitting the main GAM for each species-year using a randomly chosen training subset (90% of the data). We obtained model predictions for the remaining 10% of the data and compared model predictions with observed detections. We assessed two model performance metrics: Cohen’s kappa calculated in the PresenceAbsence package v. 1.1.9 (Freeman & Moisen 2008), and the correlation for binned detection averages. Cohen’s kappa is used to evaluate agreement for binary data, and is particularly useful when overall detection rates are low (Cohen 1960; Strimas-Mackey *et al*. 2020); kappa values near one indicate strong agreement. Our fitted GAMs had a median kappa of 0.29 (SD = 0.07, range 0.09 to 0.45; Figure S3), with a distribution that closely matches that of other recent eBird studies (e.g., mean of 0.31 in Strimas-Mackey *et al*. 2020). Additionally, we computed the correlations between predicted and observed detection rates averaged over space and time. We binned test data into 0.1° gridded coordinates and 10-day bins, and then took the average predicted and observed detection rates for any bins with at least 20 checklists. As further validation, the model predictions were strongly correlated with observed values, with a median correlation coefficient of 0.77 (SD = 0.13, range 0.31 to 0.98; Figure S3).

For each species-year, we also fit a null GAM of spring migration dynamics. The null model had the same structure as the main GAM above, but did not contain HPOP as a predictor. Hence, the null model describes spring migration dynamics without modelling an explicit effect of spatial variation in HPOP. We used the predictions of these species-year null models to estimate navigational bias, as described in the sections below. Figure 1C shows an example of the null model for the bay-breasted warbler; an additional example for the cedar waxwing is shown in Figure S2 of the supplement.

### 2.4 Estimates derived from models

We used the fitted GAMs to estimate two key metrics for each species-year: HPOP_mig_, the average level of HPOP experienced during spring migration, and HPOP_breed_, the average level experienced during breeding (Figure 1D). First, we defined the beginning and end of the spring migration phase for each species-year based on daily changes in the species’ average latitude, as estimated from the fitted GAM (see supplement for details of this procedure). Then, for each date in a given species-year analysis, we calculated the species’ daily HPOP as a weighted average, by taking the average HPOP for all sites in the species’ range, weighted by model-estimated encounter rates. We calculated HPOP_mig_ as the average of these daily values for the spring migration phase, and HPOP_breed_ as the average of these daily values for the breeding phase. An example is shown in Figure 1D for the bay-breasted warbler. Larger values of these metrics indicate that a species experiences greater human population densities during that particular phase. Further details of these estimates are provided in the supplement.

We used the rptR package v. 0.0.22 (Stoffel *et al*. 2017) to estimate repeatability, R, of HPOP_mig_ and HPOP_breed_, computed as the proportion of total variation attributed to differences among species. We included year as a categorical fixed effect when estimating R. To check for phylogenetic signal in both HPOP_mig_ and HPOP_breed_, we calculated Pagel’s lambda using the phytools package v. 0.7-80 (Revell 2012) and a phylogeny obtained from BirdTree.org (Jetz *et al*. 2012, 2014). A value of lambda near one indicates that values are highly similar among closely related species, whereas a value near zero indicates that trait variation is independent of the phylogenetic structure.

To further describe migration and breeding ecology, we examined the association between the breeding Species Specialization Index (SSI) and each of HPOP_mig_ and HPOP_breed_. Breeding SSI quantifies how specialized bird species are in terms of their breeding habitat (Julliard *et al*. 2006; Martin & Fahrig 2018; Di Cecco & Hurlbert 2022); larger SSI values indicate that a species uses a narrower range of habitat types when breeding. We examined these correlations for 59 species in our sample that had SSI values reported in previous work (Martin & Fahrig 2018; Ziolkowski *et al*. 2022).

We defined a species’ navigational bias as the difference between HPOP_mig_ and a null expectation. To estimate navigational bias, we computed HPOP_null_ as the analog to HPOP_mig_ but calculated from the null model, which did not contain HPOP as a predictor (Figure 1C). Thus, HPOP_null_ estimates the average HPOP a species experiences during its migration if we ignore HPOP *per se* when modelling migration dynamics. For each species-year, we computed navigational bias as the difference HPOP_mig_ – HPOP_null_ (Figure 1D). If HPOP_mig_ is greater than HPOP_null_, it indicates that a species is navigationally biased toward higher HPOP sites than expected during its spring migration. If HPOP_mig_ is lower than HPOP_null_, it indicates that a species is navigationally biased toward lower HPOP sites than expected.

We defined a species’ excess exposure to HPOP during spring migration as the difference HPOP_mig_ – HPOP_breed_ (Figure 1D). Larger positive values of this estimate indicate that a species experiences a greater excess of HPOP during spring migration, relative to breeding. In contrast, negative values indicate that a species experiences less HPOP during its migration than it does during the breeding phase. We report the fold-change by exponentiating this difference because HPOP_mig_ and HPOP_breed_ are expressed on a natural log scale.

### 2.5 Comparative analysis

We used a comparative analysis to investigate sources of variation in excess exposure to HPOP during migration. We fit a Bayesian phylogenetic regression model in the MCMCglmm package v. 2.32 (Hadfield 2010) (n = 189 species-year measures from 63 species). The response variable was the estimate of excess exposure to HPOP during spring migration (HPOP_mig_ – HPOP_breed_). As predictors, we considered several traits related to migration biology: the Julian date when the species starts its northward migration within North America, the overall distance the species migrates, and the time of day (nocturnal or diurnal) when its migration flight primarily occurs (Gauthreaux & Belser 2006; Van Doren *et al*. 2017; McLaren *et al*. 2018; Adams *et al*. 2021). Further details of these predictors are provided in the supplement. The model also included year as a categorical fixed effect. The random effects included species identity to account for repeated measures, as well as the phylogeny to account for shared ancestry. We ran 300,000 iterations after a burn-in period of 3,000 with a thinning interval of 500.

## 3. RESULTS

### 3.1 Species differences in HPOP estimates

HPOP_mig_ and HPOP_breed_ are highly repeatable (HPOP_mig_: R = 0.77, 95% CI = 0.67-0.85; HPOP_breed_: R = 0.93, 95% CI = 0.90-0.96; n = 189 species-year estimates from 63 species). This demonstrates that these estimates are robust and consistent for a given species across years. We did not detect significant phylogenetic signal in either trait (HPOP_mig_: lambda = 0.00, p > 99; HPOP_breed_: lambda = 0.32, p = 0.07), indicating that species differences in HPOP are not explained by shared ancestry.

As expected, HPOP_breed_ has a strong negative correlation with SSI (r = –0.49, p < 0.0001; see supplement Figure S4). This indicates that bird species with more specialized breeding habitat requirements breed in areas with lower average HPOP. By contrast, we found that a species’ HPOP_mig_ is not predicted by breeding habitat specialization, SSI (r = –0.09, p = 0.48, n = 59 species).

### 3.2 Navigational bias and excess exposure to HPOP during migration

Nearly all birds exhibit negative HPOP navigational bias (i.e., bias away from high HPOP sites during spring migration; Figure 2A). A small number of bird species were found to have positive HPOP navigational bias during migration, including the black-throated blue warbler, Nashville warbler, blackpoll warbler, dickcissel, cedar waxwing, orchard oriole, Baltimore oriole, chimney swift, and song sparrow.

**Figure 2.**
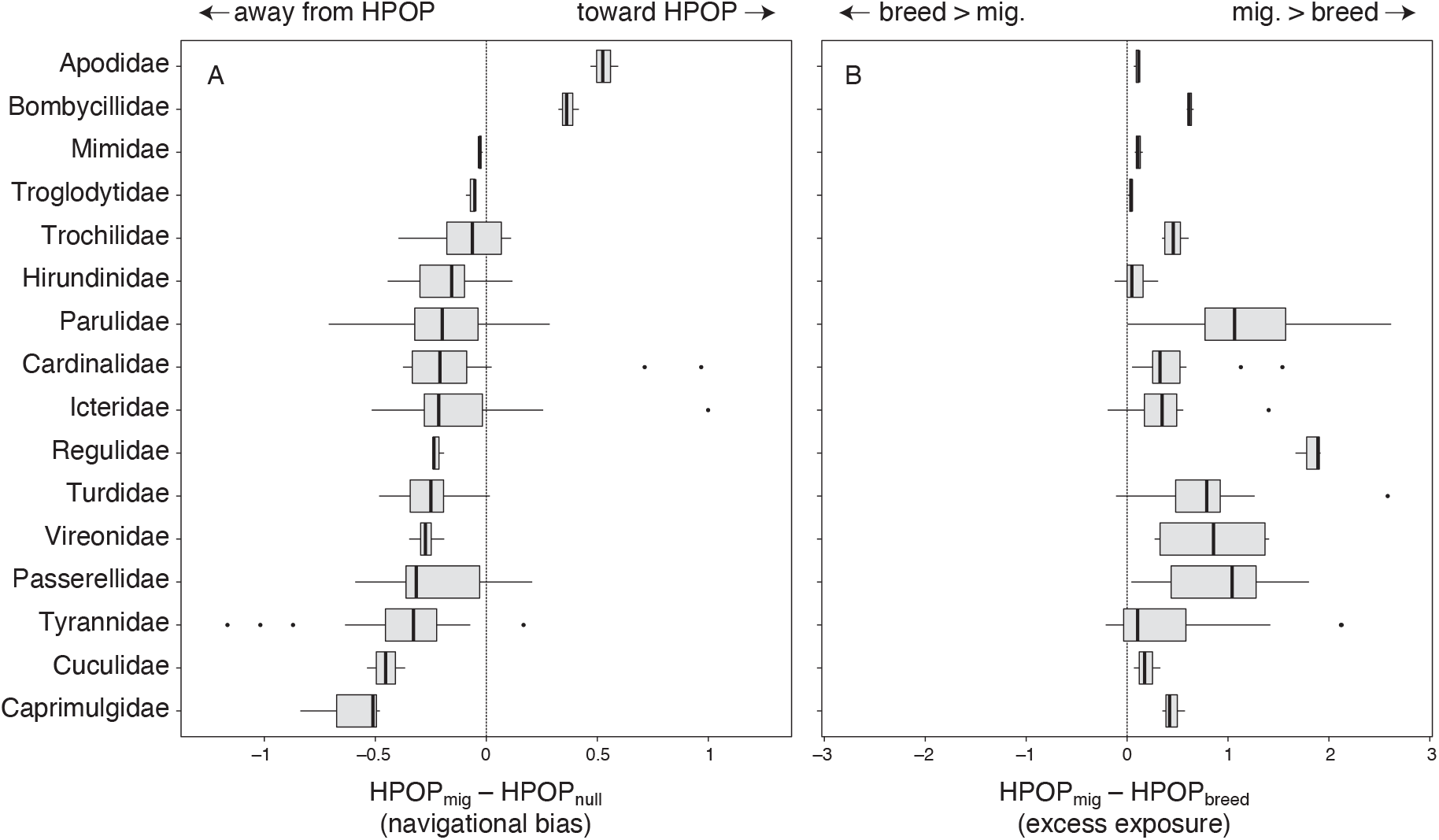
Most bird species have a negative HPOP navigational bias, yet experience much greater HPOP during migration as compared to breeding. A) Navigational bias (HPOP_mig_ – HPOP_null_) during spring migration. Most species have a negative navigational bias, indicating that they experience less HPOP during migration than expected based purely on latitudinal progress. B) Excess exposure to HPOP during spring migration (HPOP_mig_ – HPOP_breed_). Nearly all species experience far greater HPOP during migration as compared to breeding. The sample size for A-B is 189 species-years; the figure shows 16 avian families represented in this study.

Despite the widespread negative navigational bias, nearly all bird species experience much greater HPOP during spring migration than they do during breeding (Figure 2B). On average, species in our study experience 2.7-fold greater HPOP during migration as compared to breeding (range = 0.8 to 14-fold difference). The bird species with the largest excess exposure to HPOP during migration are the black-throated blue warbler, bay-breasted warbler, and Nashville warbler (see also supplement Table S1).

### 3.3 Comparative analysis

Excess exposure to HPOP during migration is predicted by three key traits (Figure 3; Table S2). The bird species with the greatest excess exposure are those that migrate primarily at night, start their migration within North America earlier in the year, and tend to migrate greater distances overall. Note that the relation with migration distance is relatively weak.

**Figure 3.**
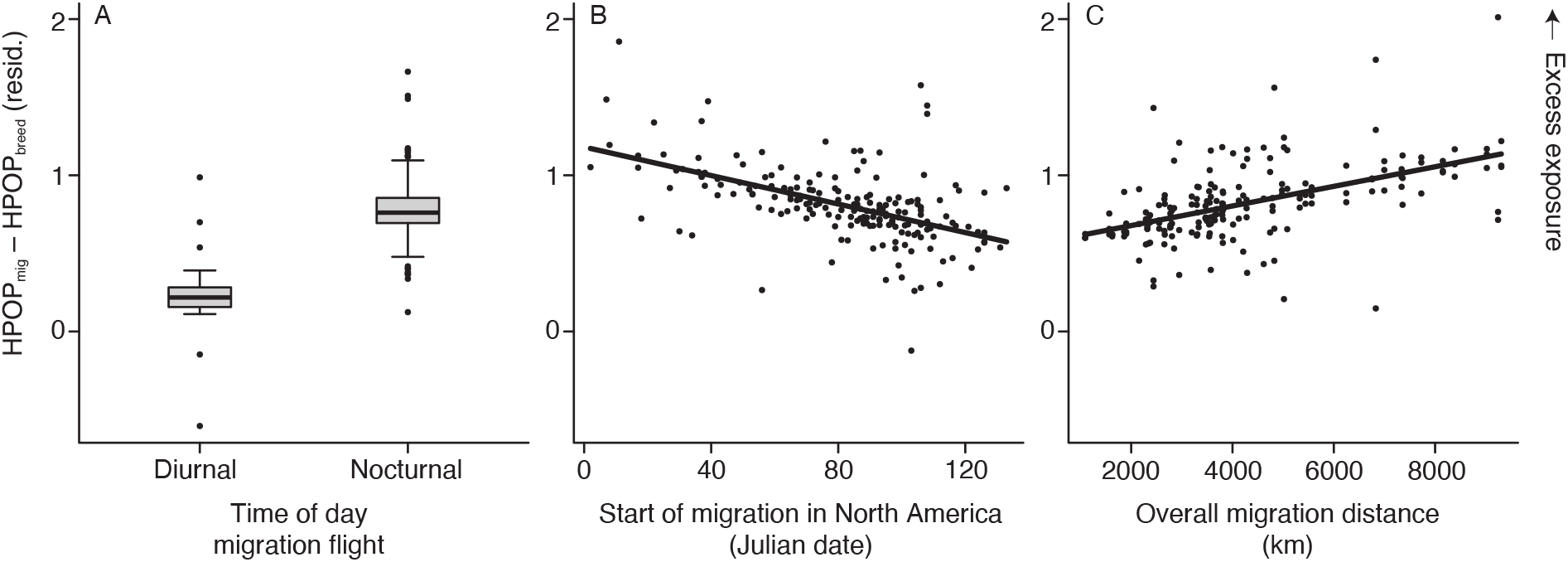
Excess exposure to HPOP during migration is associated with several key migration traits. Partial residual plots for the analysis of excess HPOP during migration (n = 189 species-years). Each panel shows the relationship between residual excess HPOP and a given predictor, after accounting for other predictors in the model. For visualization purposes, these partial residual plots were generated from a model that does not account for phylogeny. The results of this model were consistent with those of the full phylogenetic analysis. See the supplement for additional details of this analysis.

## 4. DISCUSSION

We investigated how spring migration flight of terrestrial birds is shaped by human landscape alteration across North America. Our results indicate that migrating birds are navigationally biased away from areas of high human population density, and yet they experience much greater human densities during migration than they do during breeding (Figure 2). Nevertheless, exposures to human density in North America are often elevated by 2- to 3-fold during migration, as compared to breeding (Figure 2B). Taken together, these key findings highlight how migration is a time of excess exposure to human disturbance and its associated threats (Loss *et al*. 2013, 2014, 2015; Machtans *et al*. 2013; Van Doren *et al*. 2017; Korpach *et al*. 2022).

Our results also establish that species differ repeatably in our estimates of HPOP_mig_, the average level of human density experienced by a species during migration. This indicates that some species may be more vulnerable than others to anthropogenic sources of mortality during migration (Loss *et al*. 2015). Notably, neither HPOP_mig_ nor HPOP_breed_ had detectable phylogenetic signal within the sample of terrestrial migratory birds in our study. This leads us to hypothesize that variation in species-average levels of HPOP_mig_ within North America may be explained primarily by geographic ranges and historic migration routes, rather than other evolved ecological and life history traits. HPOP_mig_ had no detectable association with breeding habitat specialization, although HPOP_breed_ was readily predicted by species differences in breeding habitat specialization (with more specialized breeders using areas with lower HPOP; Bonier *et al*. 2007; Evans *et al*. 2011; Ducatez *et al*. 2018; Callaghan *et al*. 2019; Palacio 2020; Patankar *et al*. 2021). Importantly, this finding highlights how a species’ ecological requirements during migration are not necessarily the same as during breeding (Lin *et al*. 2020). There is a major lack of information on avian ecology during migration (Faaborg *et al*. 2010). Given that birds can use very different environments even throughout migration and given the documented sources of mortality during migration (Loss *et al*. 2013, 2014, 2015; Machtans *et al*. 2013; Van Doren *et al*. 2017; Korpach *et al*. 2022), there is a crucial need for future research that maps out habitat and resource requirements for many bird species throughout the annual cycle, particularly during migration.

Although the average level of HPOP during migration was not explained by phylogeny or habitat specialization, we found that *excess* HPOP during migration (as compared to breeding) is explained by several traits (Figure 3). First, species that migrate at night are exposed to greater excess human density. Nocturnally migrating species are known to be disoriented by artificial light at night (Gauthreaux & Belser 2006; Adams *et al*. 2021), which may lead birds to spend more time in human-modified areas. During overcast conditions, some birds may also use artificial light to navigate (Weisshaupt *et al*. 2022). This may cause birds to steer towards anthropogenically modified landscapes where they are vulnerable to light disorientation effects, collisions with man-made objects, and air pollution (La Sorte *et al*. 2022b, a). For many of these species, dark-connected skies along their migration routes may be an important resource for successful navigation (Korpach *et al*. 2022).

Second, we found that species that begin migrating through North America earlier in the year are exposed to greater excess human density. Early migrants include species that can overwinter at higher latitudes (e.g., within North America), and those with broader physiological tolerances. We hypothesize that these broader tolerances allow early migrants to be more flexible in terms of habitat use and use of human-dense sites during migration (Bonier *et al*. 2007; Marzluff 2017; Ducatez *et al*. 2018; Isaksson 2018; Callaghan *et al*. 2019; Patankar *et al*. 2021). Third, we found a weak effect of overall migration distance: all else being equal, species that migrate farther tend to be exposed to greater excess human density. Longer-distance migrants have a faster overall pace of migration (La Sorte *et al*. 2013; Schmaljohann 2019) characterized by shorter and/or fewer stopovers. Owing to these challenges and faster migration pace, we hypothesize that longer distance migrants may be less able to choose stopover sites, which may lead to them stopover in urban areas more often even if the conditions of these sites are less favourable.

Migration routes and navigational responses are the product of both genetic and cultural evolution (Pulido 2007). Human development of the landscape has proceeded at a pace that far exceeds what evolution can match. Our results indicate that the majority of terrestrial migrants have a navigational bias away from human density at the population level (Figure 2A). This response could be driven by avoidance behaviours on the part of individuals (i.e., individuals avoiding cues of human-density during their migration flights and/or stopovers areas). This response could also be driven at the population level over longer time scales, via the development of migratory routes that circumvent human-dense areas when possible. It is important to note that avoidance at the population level does not preclude attraction at the individual level; these two mechanisms can co-occur even if population- and individual-level responses are opposing, and our methods do not allow us to resolve the underlying mechanisms. For example, previous work has shown that one facet of urbanization, artificial light, has both attractive and repulsive properties depending on the spatial scale under consideration (McLaren *et al*. 2018). Additionally, some bird species may continue to follow sub-optimal migration routes if they were inherited or learned from other individuals. To fully understand the impact of human landscape development on migrating birds, additional research is needed on how individual birds are responding to human density at finer scales (Bonnet-Lebrun *et al*. 2020).

Our findings have implications for avian conservation. Species such as the black-throated blue warbler, Nashville warbler, and blackpoll warbler, are both navigationally biased *toward* human-dense areas and experience far greater human density during migration as compared to their preferred breeding sites. These species may either be able to exploit resources in anthropogenically modified areas, or they may be disoriented by features in these areas, such as artificial light at night, that entrap them. Other species that have far greater excess exposure combined with a navigational bias away from human-dense areas may be forced to navigate through areas that they otherwise would have avoided; these include species such as the bay-breasted warbler, ruby-crowned kinglet, and yellow-bellied flycatcher. We have provided additional examples of this in the supplement (Figure S5). An important next step is to determine whether and when the use of human-dense sites during migration can be a threat or a benefit, both in the short and long term. Given that humans have modified areas that may be crucial for migration, billions of birds may be forced to navigate highly modified landscapes despite a preference to avoid them.

## ACKNOWLEDGEMENTS

We thank eBird and its volunteers, the Cornell Lab of Ornithology, and the Center for International Earth Science Information Network for providing data. We also thank Adam C. Smith, Christina Davy, Gabriel Blouin-Demers, Barbara Frei, Krista De Groot, Brandon Edwards, and Allison Binley for their invaluable feedback on this study.

