## supplement for "Migration and the excess exposure of birds to human density in North America"

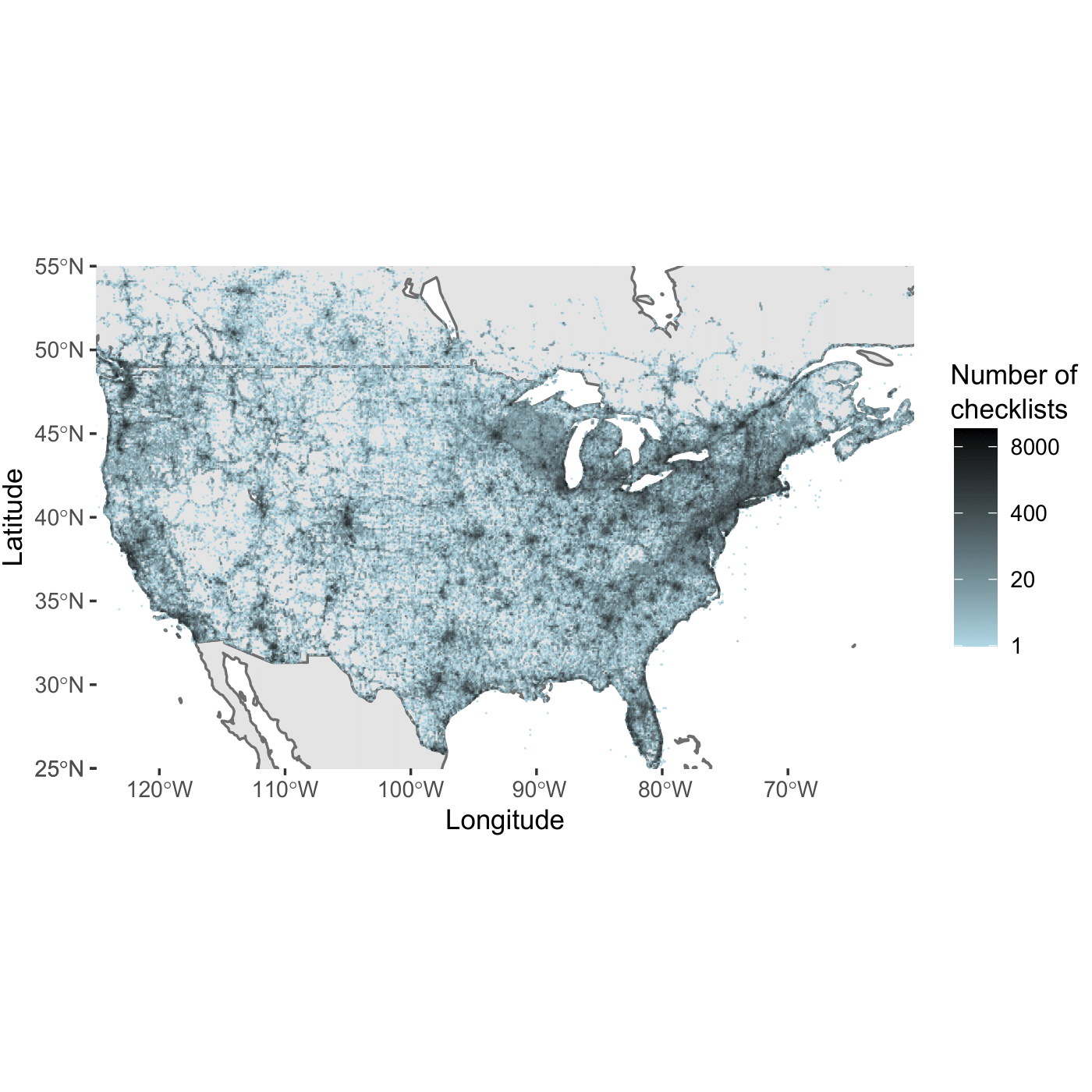

**Figure S1. Distribution of eBird checklists across the United States and Canada.** Coloured points represent eBird checklist sites in the Sampling Event Dataset gridded to the nearest 0.1°. Note that checklist frequency is on a log-scale, and the legend shows back-transformed values. For visualization purposes, checklists that occurred at latitudes above 55ºN are not shown.

**Species selection criteria**

To sample species for this study, we used three main criteria. First, we prioritized terrestrial bird species that are common and broadly-distributed throughout North America and that have breeding population sizes in the millions, based on estimates from Rosenberg et al. (2019). This first criterion was important for our ability to obtain robust estimates of migration dynamics based on eBird observations. Second, we chose species to attain a sample that included both nocturnal and diurnal migrants. Lastly, we selected species to represent a wide range of families as shown in Figure 2 of the main text.

Table S1 below provides additional information on each of the 63 study species. The majority of these species breed either cross-continentally or in the eastern/central regions of North America, with only three exclusively western breeders (the rufous hummingbird, Hammond’s flycatcher, and Townsend’s warbler).

**Table S1. Bird species sampled in this study.** Species are ordered by phylogeny (Jetz *et al.* 2012, 2014). HPOP_mig_ is the average level of HPOP experienced during the migration phase (on a natural log scale). The North American range column was determined based on range maps provided in Birds of the World (Billerman *et al.* 2022). Note that the full dataset and all estimates used in this study are available in the associated open data repository at the link provided in the main text.

| Species | Common name | Time of day of migration | HPOP_mig_ | Navigational bias | NORTH AMERICAN RANGE |
| --- | --- | --- | --- | --- | --- |
| *Selasphorus rufus* | Rufous hummingbird | Diurnal | 1.6 | -0.5 | Western |
| *Archilochus colubris* | Ruby-throated hummingbird | Diurnal | 1.1 | -0.03 | Eastern/central |
| *Chaetura pelagica* | Chimney swift | Diurnal | 0.3 | 1.6 | Eastern/central |
| *Chordeiles minor* | Common nighthawk | Diurnal | 1.4 | -1.8 | Cross-continental |
| *Myiarchus crinitus* | Great crested flycatcher | Nocturnal | 0.4 | -0.4 | Eastern/central |
| *Tyrannus tyrannus* | Eastern kingbird | Diurnal | -0.2 | -1.3 | Cross-continental |
| *Empidonax virescens* | Acadian flycatcher | Nocturnal | -0.4 | -1.4 | Eastern |
| *Empidonax minimus* | Least flycatcher | Nocturnal | 2.9 | -0.09 | Eastern/central |
| *Empidonax hammondii* | Hammond's flycatcher | Nocturnal | 0.7 | -2.4 | Western |
| *Empidonax alnorum* | Alder flycatcher | Nocturnal | 1.7 | -0.8 | Cross-continental |
| *Empidonax traillii* | Willow flycatcher | Nocturnal | 1 | -0.9 | Cross-continental |
| *Empidonax flaviventris* | Yellow-bellied flycatcher | Nocturnal | 5.7 | -1.8 | Cross-continental |
| *Contopus virens* | Eastern wood-pewee | Nocturnal | -0.3 | -1 | Eastern/central |
| *Vireo olivaceus* | Red-eyed vireo | Nocturnal | 1 | -0.7 | Cross-continental |
| *Vireo solitarius* | Blue-headed vireo | Nocturnal | 4.1 | -0.9 | Eastern/central |
| *Dumetella carolinensis* | Grey catbird | Nocturnal | 0.3 | -0.08 | Cross-continental |
| *Catharus ustulatus* | Swainson's thrush | Diurnal | 2.2 | -0.8 | Cross-continental |
| *Catharus minimus* | Grey-cheeked thrush | Nocturnal | 5.09 | -0.5 | Cross-continental |
| *Catharus fuscescens* | Veery | Nocturnal | 2.2 | -0.8 | Cross-continental |
| *Hylocichla mustelina* | Wood thrush | Nocturnal | 0.04 | -1.1 | Eastern |
| *Troglodytes aedon* | House wren | Nocturnal | 0.1 | -0.2 | Cross-continental |
| *Bombycilla cedrorum* | Cedar waxwing | Diurnal | 1.9 | 1.1 | Cross-continental |
| *Regulus calendula* | Ruby-crowned kinglet | Nocturnal | 5.5 | -0.7 | Cross-continental |
| *Icterus galbula* | Baltimore oriole | Diurnal | 0.8 | 0.2 | Eastern/central |
| *Icterus spurius* | Orchard oriole | Nocturnal | 1.08 | 0.1 | Eastern/central |
| *Dolichonyx oryzivorus* | Bobolink | Diurnal | 1.4 | -0.7 | Eastern/central |
| *Spizella arborea* | American tree sparrow | Nocturnal | 3.5 | -1.09 | Cross-continental |
| *Passerella iliaca* | Fox sparrow | Nocturnal | 4.2 | -1.4 | Cross-continental |
| *Zonotrichia albicollis* | White-throated sparrow | Nocturnal | 4.6 | -0.1 | Eastern/central |
| *Melospiza lincolnii* | Lincoln's sparrow | Nocturnal | 3.5 | -1 | Cross-continental |
| *Melospiza georgiana* | Swamp sparrow | Nocturnal | 1.3 | -0.9 | Eastern/central |
| *Melospiza melodia* | Song sparrow | Diurnal | 0.2 | 0.4 | Cross-continental |
| *Spizella pallida* | Clay-coloured sparrow | Diurnal | 1.3 | -0.8 | Central |
| *Seiurus aurocapilla* | Ovenbird | Nocturnal | 2.6 | -0.9 | Eastern central |
| *Dendroica fusca* | Blackburnian warbler | Diurnal | 3.4 | -2 | Eastern/central |
| *Dendroica castanea* | Bay-breasted warbler | Nocturnal | 6.5 | -1.09 | Eastern |
| *Dendroica striata* | Blackpoll warbler | Diurnal | 5.8 | 0.2 | Cross-continental |
| *Dendroica petechia* | Yellow warbler | Nocturnal | 0.09 | -0.4 | Cross-continental |
| *Dendroica pensylvanica* | Chestnut-sided warbler | Nocturnal | 2.3 | -1 | Eastern |
| *Parula americana* | Northern parula | Nocturnal | 2.09 | -0.08 | Eastern |
| *Dendroica caerulescens* | Black-throated blue warbler | Nocturnal | 6.6 | 0.3 | Eastern |
| *Dendroica virens* | Black-throated green warbler | Nocturnal | 3.7 | -1.3 | Eastern/central |
| *Dendroica townsendi* | Townsend's warbler | Nocturnal | 2.3 | -1.5 | Western |
| *Setophaga ruticilla* | American redstart | Nocturnal | 3 | -0.09 | Cross-continental |
| *Dendroica magnolia* | Magnolia warbler | Nocturnal | 5.5 | -0.2 | Eastern/central |
| *Wilsonia citrina* | Hooded warbler | Nocturnal | 2.4 | -1 | Eastern |
| *Wilsonia pusilla* | Wilson's warbler | Nocturnal | 2 | -0.1 | Cross-continental |
| *Mniotilta varia* | Black-and-white warbler | Nocturnal | 3.1 | -0.9 | Eastern/central |
| *Seiurus noveboracensis* | Northern waterthrush | Nocturnal | 3.4 | -0.6 | Cross-continental |
| *Oporornis philadelphia* | Mourning warbler | Nocturnal | 4.5 | -0.2 | Eastern/central |
| *Geothlypis trichas* | Common yellowthroat | Nocturnal | 0.7 | -0.7 | Cross-continental |
| *Vermivora peregrina* | Tennessee warbler | Nocturnal | 4.8 | -0.3 | Cross-continental |
| *Vermivora ruficapilla* | Nashville warbler | Nocturnal | 6.3 | 0.2 | Cross-continental |
| *Spiza americana* | Dickcissel | Nocturnal | 3.3 | 1.7 | Central |
| *Passerina cyanea* | Indigo bunting | Nocturnal | 0.6 | -0.7 | Eastern/central |
| *Pheucticus ludovicianus* | Rose-breasted grosbeak | Nocturnal | 1.2 | -0.8 | Eastern/central |
| *Piranga rubra* | Summer tanager | Nocturnal | 0.8 | -0.9 | Cross-continental |
| *Tachycineta bicolor* | Tree swallow | Diurnal | -0.09 | -1 | Cross-continental |
| *Riparia riparia* | Bank swallow | Diurnal | -0.2 | -1 | Cross-continental |
| *Progne subis* | Purple martin | Diurnal | 0.7 | -0.5 | Cross-continental |
| *Petrochelidon pyrrhonota* | Cliff swallow | Diurnal | 0.5 | -0.04 | Cross-continental |
| *Hirundo rustica* | Barn swallow | Diurnal | 0.1 | -0.3 | Cross-continental |
| *Coccyzus americanus* | Yellow-billed cuckoo | Nocturnal | 0.6 | -1.4 | Eastern/central |

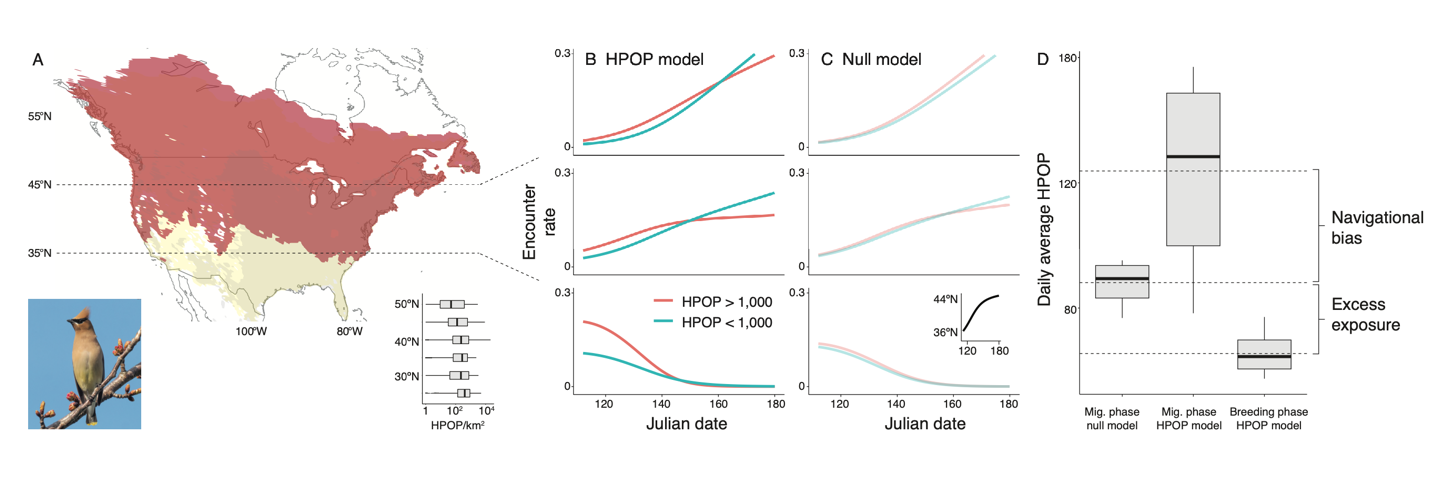

**Figure S2. Modelling spring migration dynamics for the cedar waxwing (*Bombycilla cedrorum*).** This example can be compared with the bay-breasted warbler in Figure 1 of the main text. All features of this figure follow those of Figure 1. Notably, unlike the bay-breasted warbler, the cedar waxwing is estimated to have positive navigational bias (i.e., during migration, it is overrepresented at high HPOP sites). Note that in A, the grey shaded region represents the waxwing’s overwintering areas and the yellow and orange regions represent its migration and breeding regions, respectively. Photo by Wikimedia Commons user Rhododendrites.

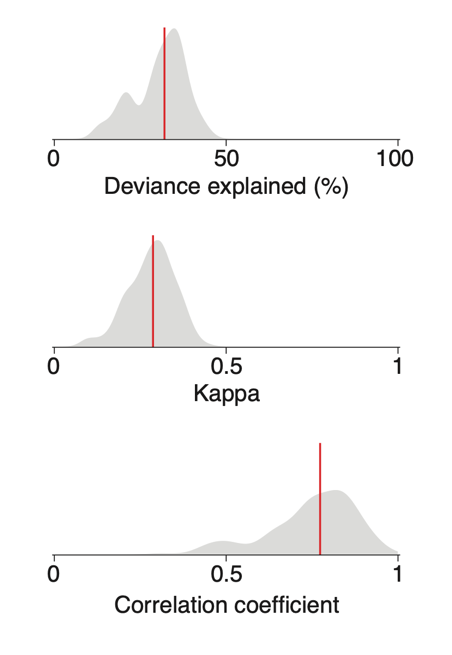

**Figure S3. Model support for migration GAMs.** Distributions for deviance explained (top), cross-validation kappa (mid), and the correlation between out-of-sample predicted and observed values (bottom) for 189 species-year models.

### **Defining the migration phase for HPOP estimates**

Migratory species vary in their overwintering ranges as well as the timing of migration and arrival in North America. To ensure that our HPOP estimates were only calculated for dates when a species was present in reasonable numbers within North America, we first defined a species as occurring in North America on the first date when the cumulative number of eBird checklist detections in North America reached 5% of the total number of checklist detections for that species-year. For any days prior, we deem the species as not yet occurring in large enough numbers to estimate encounter rates within North America.

Next, for each species-year, we used encounter rate estimates to define when the migration and breeding phases occur within North America. To define the start date of migration within North America, we examined changes in average daily latitude estimates. We calculated average daily latitudes as weighted averages. For each date, we took the mean of the latitude coordinates for all sites in the species’ range, weighted by model-estimated encounter rates. We then defined the start of migration within North America as the first date on which the species’ average latitude was at least +5 km relative to the previous day (i.e., when the average latitude for the species had moved at least 5 km northward). We defined the breeding phase as beginning when the species’ average latitude change fell below +5 km/day, or the 150^th^ day of the year (May 30), whichever date was earlier. One species, the song sparrow, never met the +5 km/day threshold for migration because it only reached daily average latitude changes of +3-4 km/day at a maximum. We therefore defined the migration phase for the song sparrow as starting on the first date when its latitudinal change was greater than +0 km/day, and ending on the last date when it met this threshold, or the 150^th^ day of the year, whichever was earlier.

The threshold of +5 km/day was chosen to represent consistent northward progress. To evaluate the robustness of estimates derived using this definition, we performed a sensitivity analysis below to test whether HPOP_mig_, HPOP_breed_, and HPOP_null_ were sensitive to changes in the threshold (Table S2). We re-calculated these metrics using three alternative thresholds: +4 km/day, +6 km/day, and +10 km/day, and examined Pearson’s correlations to compare the metric values obtained in these different threshold scenarios. The results of this analysis shown in Table S2 below demonstrate that HPOP_mig_, HPOP_breed_, and HPOP_null_ are all highly robust to changes in the migration threshold (all correlation coefficients > 0.98).

**Table S2. Robustness of HPOP estimates to the definition of the migration phase.**

| **Estimate** | **Alternative threshold (km/day)** | **Correlation with values obtained with + 5 km/day threshold** |
| --- | --- | --- |
| HPOP_mig_ | 4 | >0.99 |
|  | 6 | >0.99 |
|  | 10 | 0.98 |
| HPOP_breed_ | 4 | >0.99 |
|  | 6 | >0.99 |
|  | 10 | >0.99 |
| HPOP_null_ | 4 | >0.99 |
|  | 6 | >0.99 |
|  | 10 | 0.98 |

### **Validating HPOP estimates using eBird Status and Trends**

As an additional validation step, we compared HPOP_mig_ and HPOP_breed_ to analogous estimates derived from the eBird Status and Trends project (hereafter, ebirdst Data Products). ebirdst Data Products are publicly available estimates of species occurrence based on eBird data and using a modelling framework that also accounts for sampling biases (i.e., encounter rates). ebirdst Data Products are provided at weekly intervals and a 2.96 km spatial resolution (Fink *et al.* 2020). For each of the 63 species in our study, we used the ebirdst package v. 0.3.2 (Strimas-Mackey *et al.* 2021) to obtain weekly average encounter rates for each site in the species’ 2019 range as defined here. We used 2019 because it corresponds to one of our three sample years. For comparison, we obtained weekly average encounter rates from our fitted GAMs for 2019. We then calculated estimates of HPOP_mig_ and HPOP_breed_ from ebirdst and our comparable GAM-based procedure, for each species. Finally, we checked Pearson’s correlations for HPOP estimates derived under these two modelling frameworks. For both metrics, we found strong correlations (r = 0.84 for HPOP_mig_ and r = 0.95 for HPOP_breed_). We thus conclude that our models capture similar migration dynamics as those used in ebirdst, providing support that variation in species’ HPOP estimates are robust and generalizable.

### **Predictors for the comparative analysis**

As predictors for excess HPOP during migration, we considered several traits related to migration biology: the date when northward migration begins within North America, the overall distance that the species migrates, and the time of day (nocturnal or diurnal) when its migration flight primarily occurs.

To determine the date when the species begins its northward migration within North America, we used the first day of the year on which the species’ estimated average latitude was at least +5 km relative to the previous day, as described above.

To determine the overall distance that each species migrates, we used the ebirdst package to determine each species’ average centroid coordinates for January and June 2019 within South, Central and North America (note that we did not use our in-house GAMs, because our analysis only covered North America). These centroid coordinates were calculated as x and y coordinates weighted by ebirdst occurrence estimates. We then determined the distance between the southernmost minimum and northernmost maximum centroids, yielding an estimate of overall distance in km for each of our 63 study species. We compared these estimates against values reported by La Sorte *et al.* (2013) for 37 species that were shared in both studies and verified that the distances were highly correlated (r = 0.98, p < 0.0001).

To determine the time of day when a species migrates, we used comprehensive species accounts from the Birds of the World online resource (Billerman *et al.* 2022) to classify each species as either an exclusive nocturnal migrant (a species that migrates only at night) or a diurnal/mixed migrant (a species that migrates only during the day or during a mix of diurnal and nocturnal times). If migration timing information was not available from Birds of the World, we referred to the All About Birds website (Cornell Lab of Ornithology 2019; <https://www.allaboutbirds.org>).

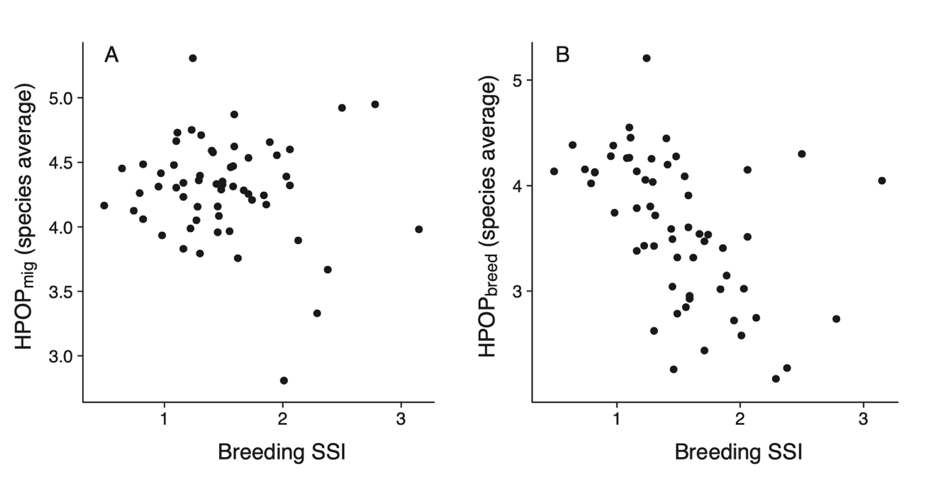

**Figure S4. Breeding SSI is negatively correlated with HPOP_breed_, but not HPOP_mig_.** Scatterplots show values for 59 species that have breeding SSI obtained from previous studies.

**Table S3. Comparative analysis of excess HPOP during migration.** Posterior means, 95% credible intervals (CI), effective sample sizes, and pMCMC values are shown. Note that parameter estimates are reported for a model where the two continuous predictors were standardized to have a mean of 0 and SD of 1.

| Predictor | Posterior mean | 95% CI | | Effective sample size | pMCMC |
| --- | --- | --- | --- | --- | --- |
| Year (2018 relative to 2017) | 0.04 | –0.06 | 0.12 | 600 | 0.37 |
| Year (2019 relative to 2017) | 0.07 | –0.03 | 0.16 | 600 | 0.13 |
| Time of day for mig. flight (nocturnal relative to diurnal/anytime) | 0.53 | 0.11 | 0.93 | 798 | 0.02 |
| Start date of NA mig. | –0.13 | –0.21 | –0.03 | 600 | 0.01 |
| Overall migration distance | 0.14 | –0.03 | 0.35 | 600 | 0.11 |

**Examples of HPOP dynamics during migration**

The average HPOP experienced by each species fluctuates over the course of its annual cycle. Based on visual inspection of these time series, we identified four general patterns that are illustrated in Figure S5 below:

1. Species with a negative HPOP navigational bias that experience excess exposure to HPOP during the migration phase as compared to breeding (e.g., the black-and-white warbler; Figure S5A; n = 12 species).
2. Species that appear to be either attracted to HPOP or navigationally neutral and that experience excess exposure to HPOP during the migration phase as compared to breeding (e.g., the American redstart; Figure S5B; n = 25 species).
3. Species with a negative HPOP navigational bias that experience similar HPOP during the migration and breeding phases (e.g., the bank swallow; Figure S5C; n = 14 species).
4. Species that appear to be either attracted to HPOP or navigationally neutral and that experience similar HPOP during the migration and breeding phases (e.g., the barn swallow; Figure S5D; n = 12 species).

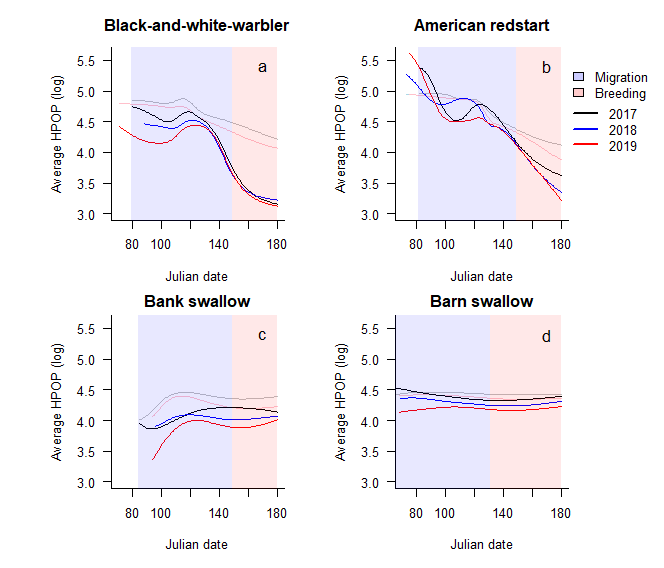

**Figure S5. Examples of HPOP dynamics**. Examples are shown for four species, the A) black-and-white warbler, B) American redstart, C) bank swallow, and D) barn swallow. In each panel, the darker lines show daily HPOP estimates. The semi-transparent lines show expectations from the null model. The migration and breeding phase are denoted by background shading.
